# Identification and genomic characterization of a novel bisegmented coronavirus in the lesser panda

**DOI:** 10.1101/2024.11.26.625429

**Authors:** Xiaoyuan Chen, Hua Guo, Niu Zhou, Zhengli Shi, Xinyuan Cui, Tianyi Dong, Mengjie Qin, Xianghui Liang, Fen Shan, Hai Wang, Qian Zuo, Shiming Peng, Wenjie You, David M. Irwin, Chan Ding, Leiping Zeng, Wu Chen, Yongyi Shen

## Abstract

Coronaviruses (CoVs) are enveloped positive-sense single-stranded RNA viruses and are renowned for their capacity to infect a diverse range of animals, including humans. In this study, we report the identification and characterization of a novel bisegmented coronavirus, designated LpCoV, from a dead lesser panda, which exhibited severe clinical manifestations and lesions in multiple organs. The LpCoV represents the first documented coronavirus with a unique bisegmented genome while maintaining the typical coronavirus morphology. One genomic segment encodes the ORF1ab polyprotein, while the other segment encodes the spike and nucleocapsid proteins, notably lacking the envelope (E) and membrane (M) genes. The pairwise patristic distances of the five concatenated domains in the replicase region (3CLpro, NiRAN, RdRP, ZBD, and HEL1) between LpCoV and α-δ coronaviruses range from 1.19 to 1.70. Based on the classification criteria established by the International Committee on Taxonomy of Viruses (ICTV), LpCoV is proposed to constitute a new genus within the Coronaviridae family. An epidemiological investigation identified five LpCoV-like viruses in lesser pandas from two different provinces (Sichuan and Jiangsu) indicating the long-term circulation and expansion of bisegmented coronaviruses in wildlife. These findings highlight the imperative for comprehensive viral surveillance in wildlife, which is essential for understanding and mitigating the risk of animal diseases and zoonotic spillover.

## Introduction

Coronaviruses (CoVs), belonging to the order *Nidovirales*, are characterized as enveloped, positive-sense single-stranded RNA viruses^1^. They are categorized into four distinct genera, alphacoronavirus, betacoronavirus, gammacoronavirus and deltacoronavirus^2^. Their genomes are similar, ranging from 27 to 32 kilobases, and encode four structural proteins: the spike (S) protein, membrane (M) protein, envelope (E) protein, and nucleocapsid (N) protein^3^. The proteins encoded by ORF1a and ORF1b are processed to produce nonstructural proteins (Nsps), which play a crucial role in viral replication^4–6^. Among the structural proteins, the M protein is the most abundant structural protein^7,8^. In contrast, the E protein is present in a relatively small amount inserted into the viral membrane, and is not indispensable for either coronavirus replication or the packaging of viral particles^9–12^.

Coronaviruses are known to infect a variety of animals, including humans, posing significant risks to both animal and public health. For example, the transmissible gastroenteritis virus (TGEV) and porcine enteric diarrhoea virus (PEDV) of alphacoronaviruses, as well as porcine deltacoronavirus (PDCoV) have caused substantial economic losses to the swine industry^13–16^. Infectious bronchitis virus (IBV), a member of the gammacoronavirus, leads to devastating economic losses in the global poultry industry^17,18^. Additionally, the feline coronavirus is capable of inducing both mild enteritis and severe systemic manifestations in cats^19^. Notably, three zoonotic coronaviruses have emerged over the past 20 years: Severe Acute Respiratory Syndrome Coronavirus (SARS-CoV)^20^, Middle East Respiratory Syndrome Coronavirus (MERS-CoV)^21^, and the more recent Severe Acute Respiratory Syndrome Coronavirus 2 (SARS-CoV-2)^22^. It’s believed that SARS-CoV and MERS-CoV originated in bats^23,24^ and subsequently transmitted to humans through market civets and dromedary camels serving as intermediate hosts, respectively^25,26^. Furthermore, SARSr-CoV-2 has been identified in both bats and pangolins^27–29^. In addition, bat-born SADS-CoV caused fatal disease in pigs^30^. These findings underscore the wildlife is the reservoirs of coronaviruses and highlight the threat of coronaviruses to public and animal health, thereby emphasizing the necessity for long-term surveillance of coronaviruses in wildlife.

In this study, a novel bisegmented coronavirus was identified in a deceased lesser panda. Phylogenetic analysis of its genome suggested that it may represent a new genus that is separate from the established alpha-, beta-, gamma-, and deltacoronaviruses. The lesser panda coronavirus (LpCoV) is the first reported and verified instance of a bisegmented coronavirus in mammals.

## Results

### Identification of a novel bisegmented coronavirus from a deceased lesser panda

On 8 April 2023, a lesser panda exhibited symptoms including reduced appetite, respiratory distress, and diarrhea, ultimately died on 12 April 2023. A necropsy revealed multi-organ damage, characterized by ascites, pulmonary hemorrhage, severe gastrointestinal distension, gastrointestinal mucosal hemorrhage, hepatomegaly, splenomegaly, and renal necrosis (refer to **Figure S1**). Histopathological examination indicated the presence of pulmonary edema, accompanied by a scattered infiltration of inflammatory cells and alveolar dilation (**Figure 1A**). There was significant coagulative necrosis observed in the renal tubules (**Figure 1B**). The hepatocytes exhibited a disorganized arrangement, accompanied by significant hepatocellular steatosis and scattered inflammatory cell infiltrates (**Figure 1C**). Additionally, there was extensive necrosis in the mucosal epithelium of the gastrointestinal tissues, along with scattered inflammatory cell infiltration (**Figure 1D**).

**Figure 1.**
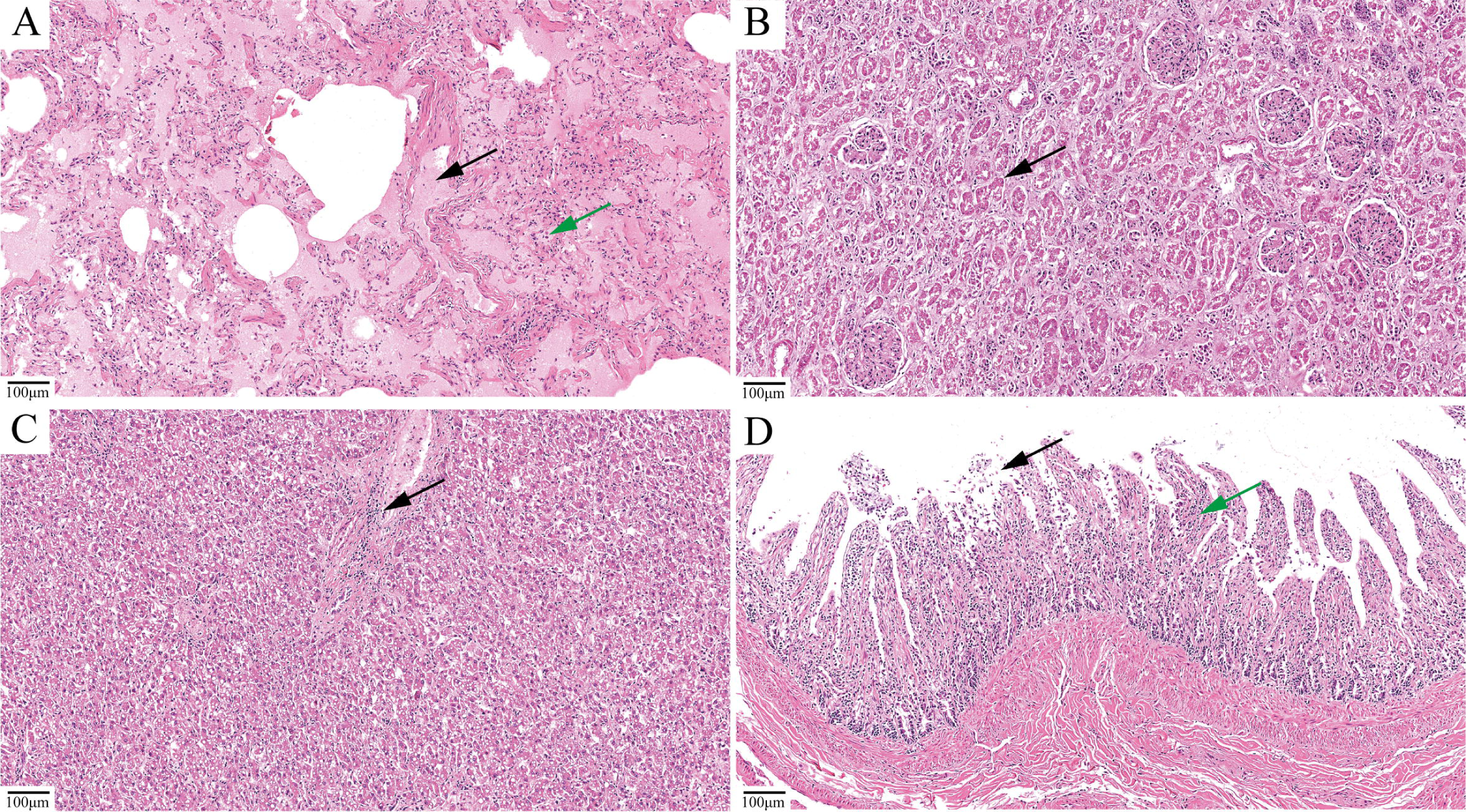
Pathological changes in the tissues of the deceased lesser panda. (A) Lung. Abnormal alveolar structure, accumulation of extravascular fluid in the alveolar lumen (indicated by black arrows) and the presence of scattered inflammatory cell infiltration (green arrows). (B) Kidney. Extensive coagulative necrosis of the renal tubular tissue (black arrows), accompanied by enlarged glomeruli and occlusion of the tubular lumen. (C) Liver. Scattered inflammatory cell infiltration (black arrows). (D) Intestine. Extensive necrosis of the mucosal glandular epithelium, detachment (black arrows), and scattered inflammatory cell infiltration (green arrows).

Using meta-transcriptome sequencing, two coronavirus segments were identified. One contig (14,249 bp) covers ORF1, including the replicase polyprotein, while the other contig (7,357 bp) contains the spike (S) and nucleocapsid (N) genes (**Figure 2A**). The average coverage for the two contigs was 22,724× fold and 12,372× fold, respectively. However, neither the E nor the M genes were detected. Both contigs exhibited poly (A) tails, indicating the presence of a bisegmented genome. Poly (A)-seq RNA sequencing, facilitated by poly (A) capture, revealed two peaks at the 3’ terminus of the two contigs (**Figure 2A**), further substantiating the hypothesis of a bisegmented genome. To eliminate the possibility of inaccurate assembly of the meta-transcriptome data, PCR was conducted to validate the integrity of the bisegmented coronavirus genome. Over-gap PCR was employed to connect the two segments of LpCoV in the anticipated direction for an unsegmented coronaviruses characterized by a typical ORF1ab-S genome. A forward primer was positioned near the 3’ end of segment 1, while a reverse primer was located near the 5’ end of the presumed segment 2 (**Figure 2B and Table S1**). However, no products were obtained, which further corroborates the notion of bisegmentation.

**Figure 2.**
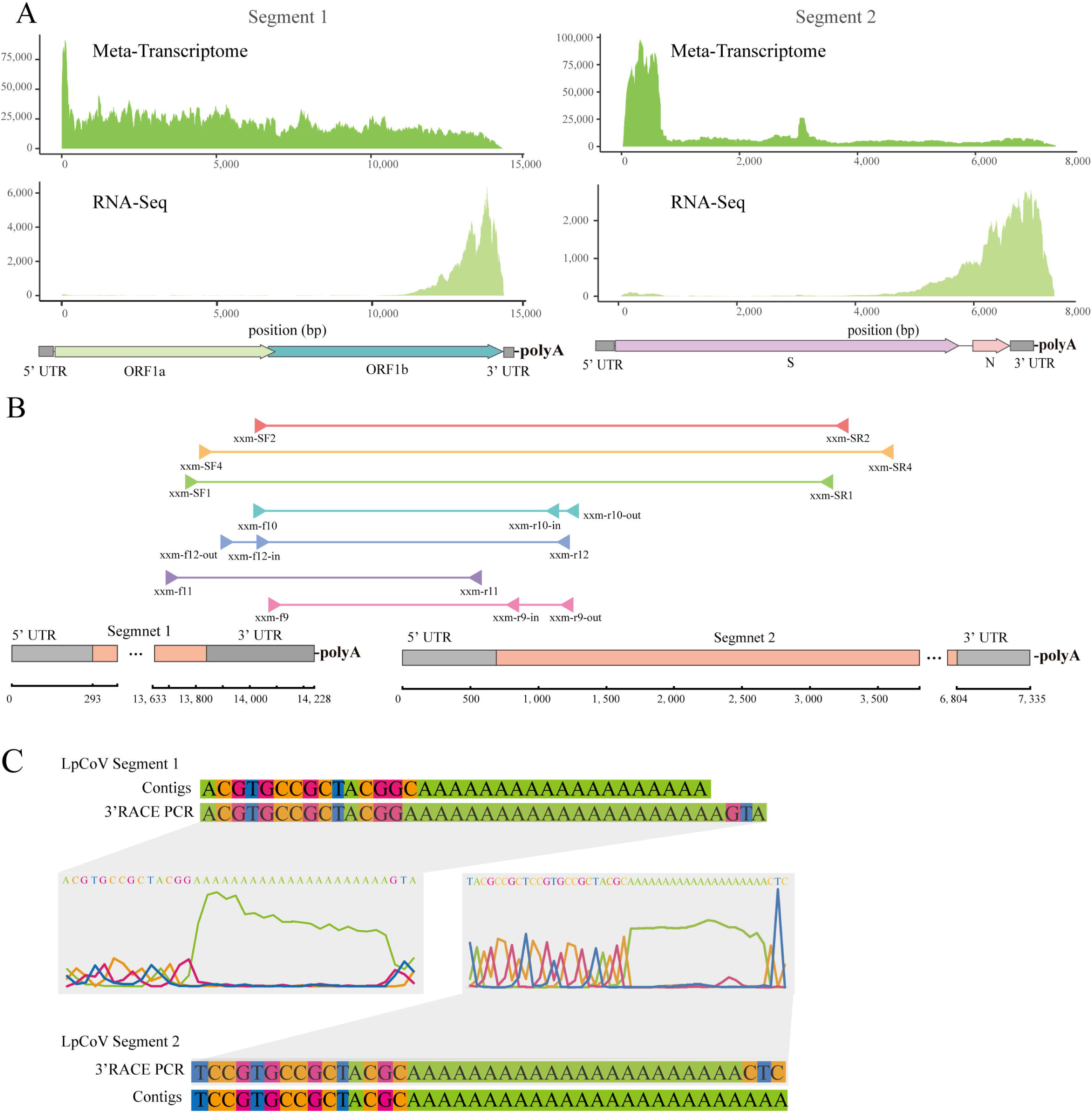
The bisegmented validation of LpCoV. (A) Sequencing depth for the two segments of LpCoV. (B) A diagram depicting the genomic locations of the nested primer pairs used for over-gap PCR. (C) 3’ RACE PCR and Sanger sequencing verified the existence of poly (A) tails in the two genomic fragments of coronaviruses.

The 5’ untranslated region (UTR) sequences for the two segments were validated by 5’ RACE PCR, and they exhibit approximately 88% nucleotide similarity (primer sequences are shown in **Table S2**). Additionally, the presence of 3’ UTR sequences and poly(A) tails in both genomic segments of LpCoV was confirmed using 3’RACE amplification (**Figure 2C**).

To avoid the omission of potential other segments, universal primers for PCR amplification were designed based on the conserved regions of the 5’/3’ UTR sequences of both segments. However, all of these attempts failed (primer sequences are shown in **Table S3).**

### Genomic characterization and phylogenetic analysis of LpCoV

The structural genes of the LpCoV genome were identified and annotated based on the established cleavage sites LXGG↓ and X-(L/I/V/F/M)-Q↓ (S/A/G) of Nsp3 and Nsp5, where X represents any amino acid and ↓ indicates the cleavage site^31^. A total of 13 nonstructural proteins, as well as the ribosomal frameshifting signal located between ORF1a and ORF1b, were identified in segment 1 (**Figure 3A**). It is noteworthy that Nsp1, Nsp2 and Nsp15 were absent from the ORF1ab region, and there is a much shorter Nsp3 (**Figure 3A**). S1 and S2 within the S proteins were characterized based on the minimal furin sequence RXXR (**Figure 3A**) ^32^. BLAST analyses in NCBI revealed that the RNA-dependent RNA polymerase (RdRp), spike, and nucleocapsid exhibit the highest protein sequence identity with the Mink coronavirus 1 (46.10%), Meles coronavirus (25.25%), and Rat coronavirus (37.50%), respectively. Therefore, representative viruses from the Nidovirales order were compiled, and a phylogenetic tree based on the amino acid sequences of the RdRp demonstrated that LpCoV did not cluster with the established alpha-, beta-, gamma-, and delta-coronaviruses, with genetic distances ranging from 0.61 to 0.72 (**Figure 3D**). Although LpCoV clustered with the Meles coronavirus, their RdRp amino acid sequences shared only 45.80% sequence identity. A multiple sequence alignment of the five concatenated domains within the replicase region (3CLpro, NiRAN, RdRp, ZBD, and HEL1) from representative coronaviruses was performed. The pairwise patristic distances (PPD) between LpCoV and Letovirinae were found to be 2.10, while distances with Pitovirinae ranged from1.90 to 2.06, and with Orthocoronavirinae from 1.19 to 1.70. According to the ICTV criteria, the threshold of PPD is set at 1.472-1.757 for genera and 2.580-3.468 for subfamilies. This suggests that the LpCoV may represent a novel genus within the Orthocoronavirinae subfamily of the Coronaviridae family, which were provisionally designated as Zetacoronavirus.

**Figure 3.**
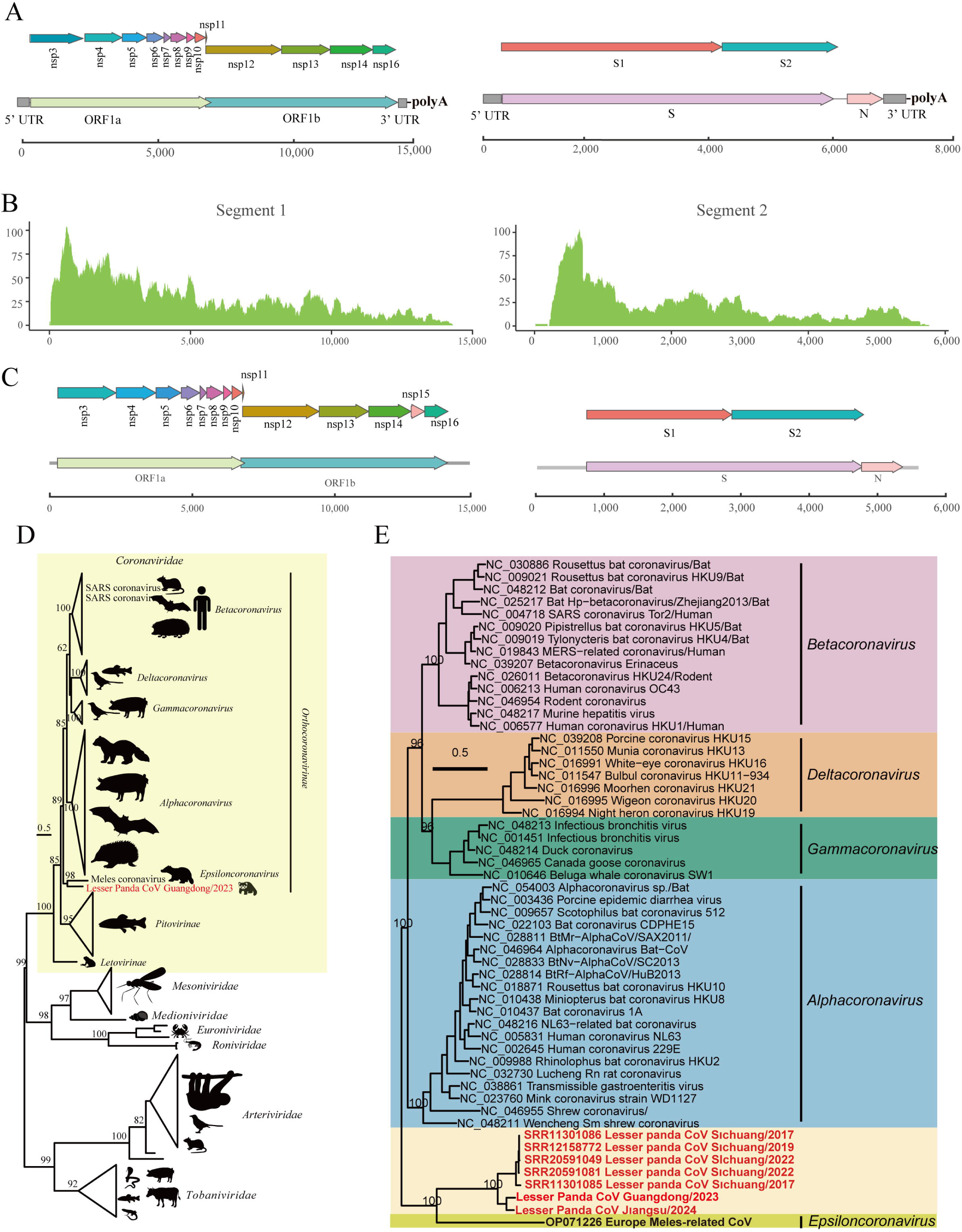
Genome characteristics and phylogenetic analysis of the LpCoV and meles coronavirus. (A) The genome architecture of LpCoV. (B) The sequencing depth for the two segments of the meles coronavirus. (C) Genome structure of the meles coronavirus. (D) A maximum likelihood phylogenetic tree based on the amino acid sequences of the RdRp gene of representative viruses with in the Nidovirale order. The LpCoV sequence is highlighted in red. Shaded colors represent the Coronaviridae. (E) A maximum likelihood phylogenetic tree based on the amino acid sequences of the RdRp gene of representative viruses of alpha-, beta-, gamma-, deltacoronavirus, and the putative Epsiloncoronavirus (Meles CoV). The LpCoV-like sequences are marked in red. The shaded colors represent different genera of Orthocoronavirinae. The scale bar indicates 0.5 amino acid substitutions per site.

In order to conduct a comparative analysis between the LpCoV and the Meles coronavirus, which has been proposed to be a newly putative genus Epsiloncoronavirus^33^, raw sequencing data for the Meles coronavirus was retrieved from the NCBI’s Sequence Read Archive (accession number PRJNA863490). *De novo* assembly and annotation were performed using our established pipelines. Similar to LpCoV, two contigs were identified for Meles coronavirus. One contig (14,429 bp) contains the replicase gene, while the other (5,767 bp) contains the spike and nucleocapsid genes, with average coverage of 19× fold and 16× fold, respectively (**Figure 3B**). The genomic architecture of the Meles coronavirus is analogous to that of the LpCoV, with the exception of an additional predicted Nsp15 within the ORF1ab region (**Figure 3C**). However, perhaps due to the low coverage, poly(A) tails were not detected in either segment.

### Epidemiological investigation of the LpCoV infection

To identify potential LpCoV infections in other zoo inhabitants, samples from 249 individuals representing 78 different species, including other lesser pandas, collected between 2017 and early 2024, were examined. These samples were organized into 126 libraries according to species, location, and health status for meta-transcriptomic sequencing (**Table S4**). The meta-transcriptomic sequencing yielded an average of 19.76 Gb of nucleotide data per library. A similar bisegmented LpCoV-like viral sequences were detected in another dead lesser panda collected from another zoo of Jiangsu Province (**Figure 3E and Figure S2A**). The replicase polyprotein, the spike, and the nucleocapsid proteins of this virus exhibited 82.8%, 59.7% and 81.2% amino acid sequence identity with LpCoV, respectively.

Furthermore, all available high-throughput sequencing data pertaining to lesser pandas from the NCBI database were gathered, which included of 40 Sequence Read Archive (SRA) accessions. LpCoV-like sequences were detected in five transcriptome datasets from lesser pandas collected in Sichuan Province, China (accession numbers SRR11301086, SRR11301085, SRR12158772, SRR20591049 and SRR20591081).

Phylogenetic analysis based on the RdRp gene revealed that these CoVs from Sichuan lesser pandas clustered with LpCoV, exhibiting an amino acid sequence identity ranging from 76.27% to 79.10% (**Figure 3E**). Among these sequences, one virus (SRR11301085 Red panda CoV 2017) was capable of being reconstructed into a full-length viral genome sequence, displaying a bisegmented genomic structure similar to that of the LpCoV (**Figure S2B**). The replicase polyprotein, the spike, and the nucleocapsid proteins of this virus exhibited 66.4%, 41.5% and 66.2% amino acid sequence identity with LpCoV, respectively.

### Tissue tropism, receptor specificity and morphology of LpCoV

RT-qPCR was employed to evaluate the tissue tropism of LpCoV across eight tissues: lung, kidney, rectum, intestine, liver, spleen, stomach, and heart. Using primers targeting the ORF1ab and spike gene, LpCoV was detected in all examined tissues except the spleen, with the highest levels of viral RNA observed in the lung, followed by the intestine, rectum, stomach, liver, heart, and kidney (**Figure 4A**). Notably, the primers targeting ORF1ab yielded viral genome copies approximately tenfold greater than those obtained with the spike primers, consistent with the meta-transcriptome data showing a higher read count for segment 1 compared to segment 2. The attempts to isolate LpCoV with the use of HEK293T, Huh7, and Vero E6 cell lines were unsuccessful. To assess the potential of LpCoV to infect host cells via interaction with ACE2, DPP4, or APN of lesser panda, a pseudotyped vesicular stomatitis virus (VSV) incorporating the spike (S) protein of LpCoV was employed to transduce HEK293T cells that express LpACE2, LpDPP4, and LpAPN, respectively. SARS-CoV pseudovirus was utilized as a positive control. The results indicated that neither the LpCoV pseudovirus nor the SARS-CoV pseudovirus was capable of transducing HEK293T cells. Notably, the SARS-CoV pseudovirus successfully transduced both HEK293T-hACE2 and HEK293T-LpACE2 cells. In contrast, the LpCoV pseudovirus did not facilitate transduction in HEK293T cells through any of the three known receptors (**Figure 4B**).

**Figure 4.**
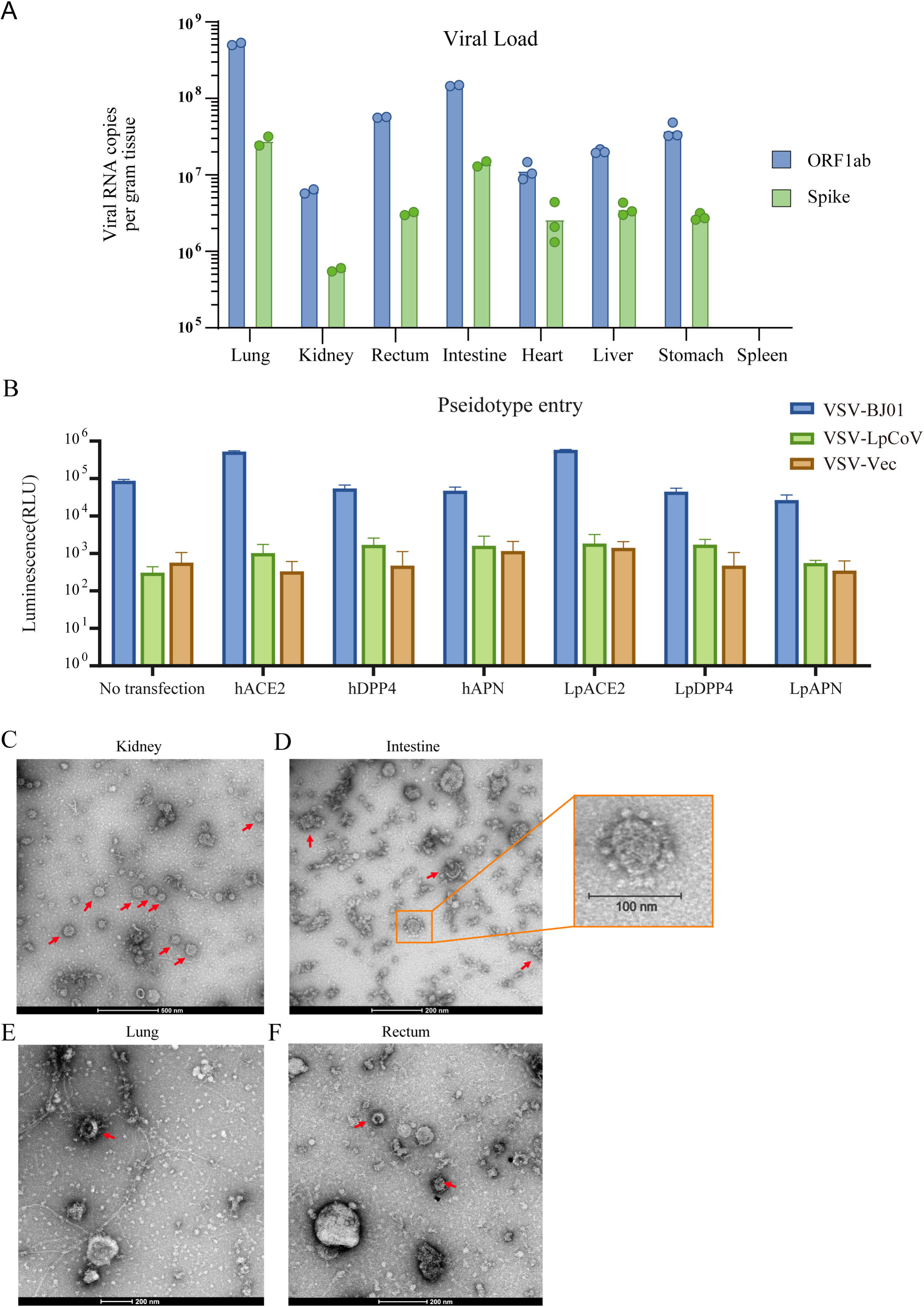
Tissue tropism and virion morphology of LpCoV. (**A**) The RNA copies of viral segment 1 and 2 per gram of tissues were quantified utilizing primer pairs that specifically target the ORF 1ab and spike genes, respectively. (**B**) Entry of VSV particles pseudotyped with LpCoVS or SARS-CoV S in HEK293T cells transiently transfected with human and lesser panda receptors. RLUs, relative luciferase units. (**C-F**) Transmission electron microscopy of purified particles of homogenized samples of the kidney, intestine, lung, and rectum tissues.

Since the *in vitro* isolation and purification of the virus was unsuccessful, to examine virion morphology, transmission electron microscopy was conducted on particles purified from homogenized lung, rectum, intestine, and kidney tissue samples via either ultracentrifugation with a sucrose cushion or density gradient ultracentrifugation. The purified particles from these tissues exhibited typical coronavirus-like morphology, with virions measuring 80-120 nm in diameter and characterized by a distinctive corona-like ring of glycoproteins on their surface (**Figure 4C-F**).

### Validation of the subgenomic mRNA expression and the transcriptional activity of LpCoV UTRs

The presence of subgenomic mRNAs is regarded as strong evidence of coronavirus replication in the infected cells. To verify the subgenome of LpCoV, CORSID was employed to predict precursor sequences^34^. Based on the transcription start site (TRS) and the transcription mechanism of coronaviruses, a potential subgenomic mRNA for LpCoV, which include a potential ORF4, is predicted (**Figure 5A**). The presence of the predicted ORF4 was assessed by probing the subgenomic mRNA with a comprehensive set of primers. As shown in **Figure 5C**, agarose gel electrophoresis revealed only a single band, and the sequencing results confirmed the presence of the N gene subgenome while absence of amplification products for ORF4 subgenomic RNA, thereby demonstrating the presence of the N gene subgenome and negating the existence of ORF4 (**Figure 5C**). In addition to an identical 5’ leader sequence, the same 3’-end structure was shared by subgenomic mRNA with the genome. (**Figure 5B and C**).

**Figure 5.**
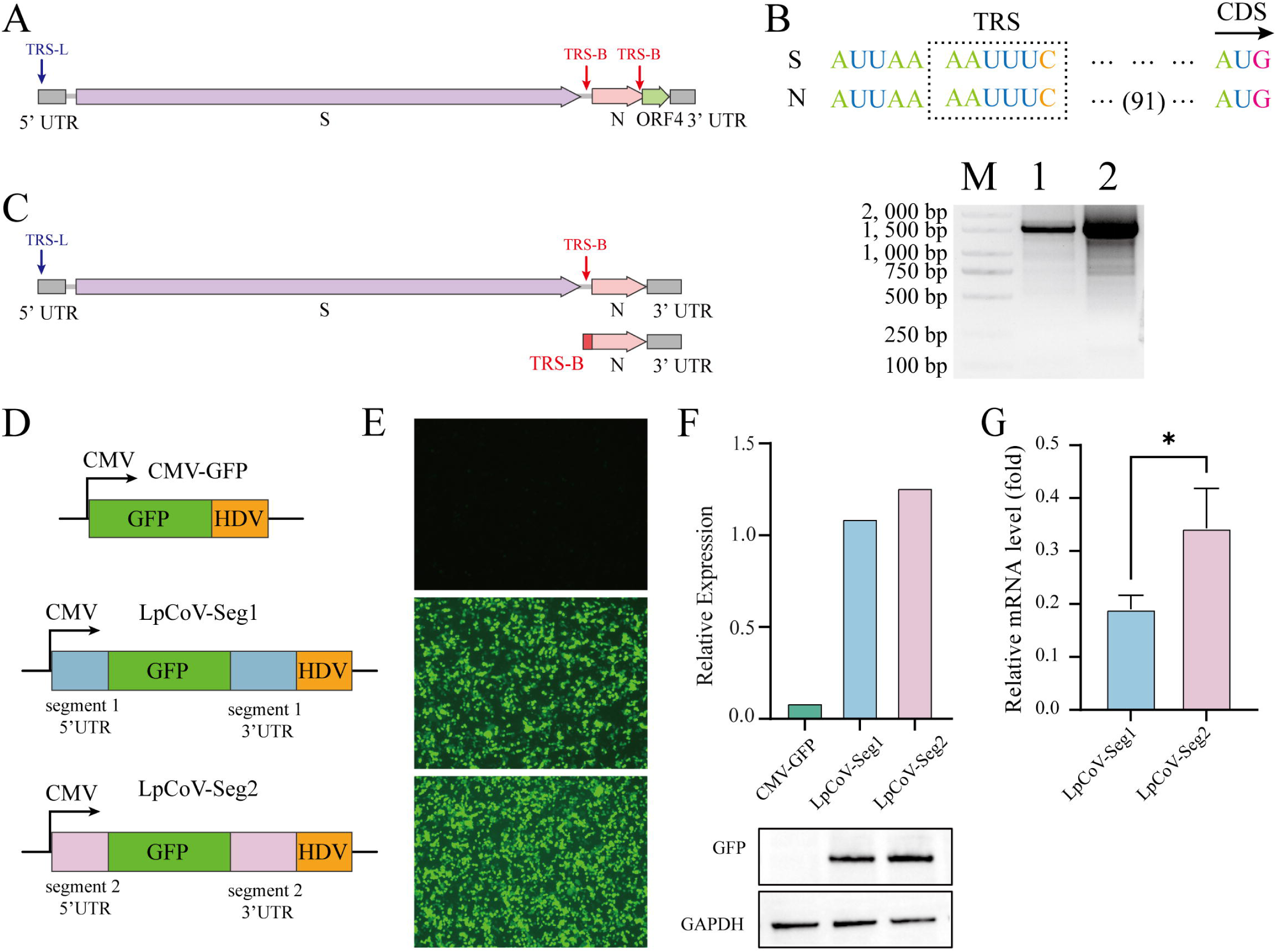
Validation of the subgenomes and the expression levels of 5’UTR of the bisegmented genome of LpCoV. (A) Schematic structures of the proposed transcribed subgenomic mRNAs. The predicted ORF4 subgenome is highlighted in a green rectangle. (B) Validation of the subgenomic and TRS sequence of LpCoV segment 2. The TRSs and CDS are indicated. The lengths of the intergenic sequences are shown with numbers. (C) Schematic structures of the transcribed subgenomic mRNAs and agarose gel electrophoresis of the PCR products of subgenomic mRNA. The numbers 1 and 2 refer to the products obtained from the first and second rounds of nested PCR for sub-genomic validation, respectively. (D) Diagrams illustrating the construction of the recombinant 5’ UTR-GFP plasmids for segment 1 and segment 2, respectively. (E) The expression of GFP was observed under a microscope. (F) Expression of the GFP protein. Cells were lysed 48 hours post-transfection and analyzed by immunoblotting with specific antibodies against GFP, using GAPDH as control. (G) The relative transcript levels of GFP were quantified using RT-qPCR. For the relative transcription levels between the 5’UTR of segment 1 and the segment 2 was calculated using the CMV-GFP control plasmid as a benchmark.

The 5’ UTR sequence is a major determinant of translation efficiency^35^. The 5’ UTRs of the two segments of LpCoV display 88% nucleotide identify. To verify the accuracy of the UTRs and to evaluate differences in expression levels between the two 5’ UTRs, their transcriptional capacities were tested. The 5’ UTR from both segments of the LpCoV genome (755 nt from segment 1 and 681 nt from segment 2) were individually cloned upstream of a reporter gene (GFP gene) (**Figure 5D**). A plasmid containing only the CMV promoter and the GFP gene was used as a control (**Figure 5D**). Each experimental sample was transfected with 2 μg of the respective plasmid. After 48 hours of transfection, cells transfected with the 5’ UTRs from both segment 1 and segment 2 plasmids displayed significant GFP protein expression (**Figure 5E**). Notably, there was no significant difference in GFP expression level between the two 5’ UTRs (**Figure 5F**).

## Discussion

Wildlife is a reservoir for emerging viruses. Decoding wildlife viruses is crucial in mitigating potential pandemic threats. Coronaviruses pose great threat to both animal and public health. Surveillance efforts have increasingly identified a series of novel coronaviruses in various wildlife and domestic species^36^. In this study, we report the identification of a novel coronavirus in a deceased lesser panda. The PPD between LpCoV and members of Orthocoronavirinae ranged from 1.19 to 1.70. Based on the ICTV criteria, it is assigned as a new genus of coronavirus, designated as Zetacoronavirus. Both the pathological anatomy and histopathological examinations indicated that multiple organ damage occurred in the deceased animal, corroborated by RT-qPCR results that confirmed viral infection in these same tissues.

The LpCoV has a bisegmented genome (**Figure 2**). A segmented genome is a common genome organization found in RNA viruses^37^, however, coronaviruses were traditionally believed to have only a single positive-sense single-stranded RNA. To our knowledge, LpCoV is the first mammalian coronavirus identified as bisegmented, thereby challenging existing paradigms regarding coronavirus diversity. Similarly, flaviviruses were initially thought to be unsegmented positive-sense RNA viruses, yet the discovery of flaviviruses with segmented genomes has significantly altered this perspective^38,39^. Coronaviruses utilize a discontinuous transcription mechanism that produces a substantial quantity of subgenomic RNAs, to avoid an early interruption of the transcription process to ensure the maintenance of their larger genomes^40,41^. These subgenomes may be encapsulated into virus particles^42,43^, potentially promoting the formation of a segmented coronaviruses. One advantage of virus genome segmentation may be manifested in better gene regulation compared to unsegmented viruses^38,44^. The segmentation pattern observed in LpCoV, which separates the S gene from the ORF1 gene, would be advantageous for the discontinuous transcription pattern of coronaviruses. Shorter genome segments may facilitate the efficient replication of segmented viruses^45^ or contribute to the stability of viral particles^46^. The gene segmentation of coronaviruses could facilitate their ability to exchange genomic segments with other viral strains. This process, known as reassortment, can lead to the emergence of novel viral strains, and is frequently documented in influenza viruses^37,47^. In this regard, a segmented genome may confer an evolutionary advantage to the virus, facilitating more efficient recombination and diversification, particularly in terms of adapting to novel hosts or environmental challenges^48,49^.

LpCoV exhibits a genomic architecture that is different from other unsegmented coronaviruses and segmented coronaviruses of aquatic vertebrates^50^. ORF1, encoded by segment 1, is notable for the absence of three non-structural proteins, Nsp1, Nsp2 and Nsp15, and features a much shorter Nsp3. Segment 2, encoding the S and N proteins, is deficient in two structural proteins, M and E. The absence of the E and M protein does not appear to affect the typical corona-like virion shape (**Figure 4C-F**). It’s established that the E protein is not essential for the replication of coronaviruses and the packaging of viral particles^10–12^, thus the absence of the E gene in this virus is not unexpected. Conversely, the M protein is required for the packaging of coronaviruses^3,7,8^. Notably, the S protein of LpCoV is larger than that of other coronaviruses, which may allow it to perform additional structural functions during the assembly process, potentially compensating for the absence of the E and M proteins. This potential compensatory mechanism warrants further investigation.

Our efforts to isolate LpCoV were not successful. In order to examine the receptor utilization of LpCoV, pseudotyped LpCoV was employed to infect HEK293T cells that were individually transfected to express ACE2, APN, and DPP4. While the positive control demonstrated successful infection with SARS CoV pseudoviruses, no indications of increased infection or viral entry were detected for LpCoV. This finding implies that none of the aforementioned receptors serve as entry receptors for LpCoV (Figure 4B).

To assess the prevalence of this virus in nature, we conducted an epidemiological study based on meta-transcriptome of 249 zoo animals, representing 78 species, alongside all available high-throughput sequencing data for lesser pandas from the NCBI database. These analyses identified the presence of LpCoV-like viruses in lesser panda samples from Jiangsu and Sichuan Provinces, China, collected between 2017 and 2024 (**Figure 3C**). The genetic similarity among these viruses was very low, indicating a widespread distribution and long-term circulation of a bisegmented coronaviruses in wildlife. Given its lethal in a lesser panda, increased surveillance is imperative to estimate the potential threat posed by this virus to this endangered species.

In conclusion, we characterized a novel bisegmented coronavirus, LpCoV, from a deceased lesser panda. Furthermore, we also identified LpCoV-like viruses in other lesser pandas through the analysis of available high-throughput sequencing data. Phylogenetic analysis suggests that the LpCoV may warrant classification in a new genus, separate from the established alpha-, beta-, gamma-, and deltacoronaviruses. This discovery not only enhances the diversity and classification of coronaviruses, but also underscores the necessity for increased vigilance regarding novel viruses in wildlife.

## Methods

### Sample collection

During a necropsy conducted on April 12, 2023, at a zoo in Guangdong province, China, tissue samples from a deceased lesser panda were obtained, including specimens from the lung, heart, liver, spleen, kidney, intestine, and stomach. A portion of these tissue samples was preserved at −80°C for future analysis, while the remainder was fixed in 4% buffered formalin for histological examination. Additionally, samples from 249 animals, encompassing 78 different species, including other lesser pandas housed in the zoo, were collected between 2017 and early 2024 to assess potential infection by LpCoV.

### Metatranscriptome sequencing and viral genome assembly

Total RNA was extracted using TRIzol (Invitrogen). The quantified RNA was subsequently used for library construction using the NEBNext Ultra Directional RNA Library Prep Kit and sequenced on the NovaSeq 6000 platform (PE150, Illumina), yielding approximately 19.76 Gb of paired-end reads generated per sample, with the exception of the deceased lesser panda, for which 57 Gb of data was generated. To reduce index hoping, the strategy “prepare dual indexed libraries with unique indexes” was implemented, and unique dual indexes (UDI) were attached to the ends of the libraries for cross verifying the indexes. In addition to the metatranscriptome sequencing, RNA-seq for enriched RNA with 3’ polyadenylated tails was conducted for the deceased lesser panda, and was used to enhance the quality of the viral genome. Low-quality raw reads were filtered using Fastp^51^. Since assembling after filtering host reads may lose some data, all clean reads were *de novo* assembled using MEGAHIT^52^. Target contigs were identified using Blastn (v2.12.0+) searches against the Coronaviridae family in the non-redundant nucleotide database. To assess coverage and depth, thereby ensuring the quality of the assembly, clean reads were remapped to the assembled contigs using Bowtie2^53^ and Samtools^54^. To further enhance the quality of the viral genome, SAM files corresponding to the reads mapping to the viral genomes were manually examined with Geneious^55^, with efforts made to extended the ends as much as possible. Open reading frames (ORFs) in the viral genome were annotated by Geneious^55^.

### Phylogenetic analysis

To elucidate phylogenetic relationships, reference strains from the Coronaviridae family were downloaded from the NCBI GenBank database. Multiple sequence alignments were generated with a high-speed and iterative refinement method (FFT-NS-i) implemented in MAFFT^56^. The sequence alignments were subsequently refined using MEGA7^57^ and trimAI^58^. Maximum likelihood (ML) trees were constructed by retrieving the best-fit model of nucleotide or amino acid substitution using IQ-TREE^59^ employing 1000 nonparametric bootstrap replicates. The phylogenetic trees were visualized using Dendroscope^60^.

Taxonomic classification was determined based on the distance matrix and ICTV standards (https://ictv.global/report/chapter/coronaviridae/coronaviridae). A concatenated multiple amino acid alignment, encompassing the domains 3CLpro, NiRAN, RdRP, ZBD, and HEL1 from coronaviruses, was generated using Clustal Omega ^61^ and subsequently trimmed with TrimAl^58^. This alignment was then used to construct a maximum likelihood tree with IQ-TREE, while pairwise patristic distances were calculated following the methodology outlined by Zhang et al^62^.

### RNA extraction and whole genome amplification

Viral RNA was extracted from 140 μL of tissue milling supernatant using the QIAamp Viral RNA Mini Kit (Qiagen, Germany). RNA was subsequently eluted in 60 μL of AVE buffer. 3 μL of the eluted RNA was used as a template for the 5’ and 3’ rapid amplification of cDNA ends (RACE) immediately. Gaps in the genome were filled by PCR amplification of the corresponding regions using specific primers. Complete genome sequences were verified via Sanger sequencing of the fragments amplified with a comprehensive set of primers designed to encompass the entire genome. The 5’/3’ RACE procedure was conducted to ascertain the terminal sequences of the genomes using the HiScript-TS 5’/3’ RACE Kit (Vazyme, Nanjing, China) according to the manufacturer’s instructions.

### Subgenome identification and sequencing

Nested subgenomic mRNAs were generated during the replication cycle of non-segmented coronaviruses. To investigate whether the identified bisegmented coronaviruses maintain this characteristic of coronaviruses, primers were designed to detect the presence of viral subgenomic mRNAs in samples confirmed to be positive for the LpCoV. The amplicons, which exhibited specific lengths corresponding to the anticipated sizes, were purified and then cloned into pTOPO vectors for sequencing (primer sequences are shown in **Table S5**). Additionally, other suspicious bands observed on the agarose gels were also excised, purified, and then sequenced directly.

### Histopathological examination

Histological analyses were performed on various tissues from the deceased lesser panda, including lung, heart, liver, spleen, kidney, intestine, and stomach. The collected tissues were cut into small pieces and subsequently fixed in 4% buffered formalin for 48 hours. Following fixation, the tissues were embedded in paraffin, sectioned at a thickness of 5 μm, and stained with haematoxylin and eosin. Results were visualized using an optical microscope (Olympus, Japan).

### Virus isolation

Tissues from the lung, kidney, rectum, and intestine of the deceased lesser panda were homogenized and used to inoculate HEK293T, Huh7, and VeroE6 cell lines for purpose of virus isolation in a biosafety level 2 laboratory. These cell lines were confirmed to be free of mycoplasma contamination using the LookOut Mycoplasma PCR Detection Kit (SIGMA), and were authenticated via microscopic morphological evaluation. Cultured cell monolayers were maintained in Dulbecco’s Modified Eagle Medium (DMEM). The inoculum was prepared by grinding the lung tissue in liquid nitrogen, diluting it 1:2 with DMEM, filtering it through a 0.45-μm filter (Merck Millipore), and treating it with 16 μg/ml trypsin solution. After incubation at 37 °C for 1 h, the inoculum was removed from the culture and replaced with fresh culture medium. The cells were incubated at 37 °C and observed daily for cytopathic effects.

### Production of VSV Pseudotyped Viruses

HEK293T cells were cultured in DMEM containing 10% fetal bovine serum (FBS) and 1% Penicillin-Streptomycin (PenStrep) and were plated in 10 cm dishes coated with poly-D-lysine. These cells were transfected with 30 μg of plasmids encoding the Spike protein. The following day, the cells were washed with DMEM and subsequently transduced with VSV-ΔG-luc. After a 2-hour incubation, the viral inoculum was removed, and the cells were washed again with DMEM. Subsequently, DMEM supplemented with an anti-VSV-G antibody (Kerafast) was added to reduce the background from the parental virus. The pseudoviruses were collected 24 hours post-inoculation, filtered through a 0.45 μm membrane, and then aliquoted and stored at −80°C.

### Transmission electron microscopy

Tissues from the lung, intestine, rectum, and kidney tissues of the deceased lesser panda were minced and homogenized in DMEM with 2x Penicillin/Streptomycin medium on ice. After centrifugation at 10,000 x g for 10 minutes at 4°C and filtration through a 0.45 μm filter, the supernatants were loaded onto 1 mL of 30% sucrose in PBS buffer and centrifuged at 25,000 rpm at 4°C for 3 hours using the 90AT rotor (Beckman). The pelleted viral particles were resuspended in 100 μL of PBS and fixed with formaldehyde (working concentration 0.1%) at 4°C overnight. A 5 μL aliquot of the fixed particles was retained, and the remainder was then loaded onto a 20%-60% sucrose cushion and centrifuged at 25,000 rpm at 4°C for another 4 hours using the SW41 rotor (Beckman). The initial 5 μL sample and each sucrose gradient layer were then absorbed, stained with uranium, and examined using a transmission electron microscope (Thermo Fisher, Talos L120C) at 120 kV.

### Real-time quantitative RT-PCR (RT-qPCR)

The viral loads in various tissues of the deceased lesser panda were analyzed by RT-qPCR. In brief, 1 gram of each tissue sample was minced and homogenized. Subsequently, 50 μL of the supernatant was used for RNA extraction using a Virus DNA/RNA Extraction Kit (Vazyme, Nanjing, China), followed by RT–PCR. The primers designed for the ORF1ab were as follow: 5’-ATGTAGTCCCCACCATAACGC-3’(forward), and 5’-TTGGTGGTGCCAATAACGACA-3’ (reverse). Primers target spike gene were: 5’-GAACGGTTCCATGTTCGTGC-3’(forward), and 5’-ACTGTCTGTACAGGGTCCGT-3’ (reverse).

The transcript levels of GFP were evaluated by RT-qPCR. Total RNA was extracted from cells using the Total RNA Isolation Kit (RC112, Vazyme, Nanjing, China). The primers for GFP were: 5’-TGCCCGACAACCACTACCTG-3’ (forward), and 5’-CATGTGATCGCGCTTCTCGTT-3’ (reverse). The primers for the GAPDH gene were: 5’-GACAAGCTTCCCGTTCTCAG-3’ (forward), and 5’-GAGTCAACGGATTTGGTGGT-3’ (reverse). RT-qPCR was conducted by the HiScript® II One Step qRT-PCR SYBR Green Kit (Q221-01, Vazyme, Nanjing, China) using the C1000 Touch Thermal Cycler/CFX96 Real-time system (Bio-Rad) and the data were analyzed using Bio-Rad CFX Maestro software (version 4.1.2433.1219). Two independent runs for each sample were done.

### Western blotting

HEK293T cells were cultured in 24-well plates until they reached a confluence of 80%. Subsequently, the cells were transfected with the validation plasmid and control plasmid, respectively. Following a 48h-hour incubation, the cells were lysed using RIPA buffer containing cocktail of protease inhibitors. The resulting samples were separated by SDS-PAGE and subsequently transferred to PVDF membranes. For blocking, the membranes were incubated in PBS containing 5% skim-milk for 2 hours at 37C, followed by overnight probing with a primary antibody at 4°C. The membranes were then washed three times with washing buffer (TBST) and incubated with the secondary antibodies in TBST for 2 hours at 37C. After three washes, the secondary antibodies were detected using the ECL Western blot reagent (Vazyme, Nanjing, China). Quantification of the bands was performed using the Tanon 5200 imaging system (Tanon, Shanghai, China).

## Data availability

Sequence reads generated in this study are available from the NCBI Sequence Read Archive (SRA) database under BioProject accession number PRJNA1178809.

## Author contributions

Y.S., W.C, C.D., and Z.S. designed and supervised the research. X.Chen and W. Y. performed the validation of bisegmented genome. H.G., L.Z., T.D., and M.T. performed the virus isolation and transmission electron microscopy. N. Z., W.C., F. S., and S. P. did the necropsy and histopathological examination and sample collection. X.Cui, X.L., Q.Z., and H.W. performed the genome assembly, sequence annotation and phylogenetic analyses. Y.S., and X.Chen drafted the paper. Y.S., C.D., W.C, D.M.I., and Z.S. revised the paper. All authors reviewed and edited the paper.

## Acknowledgments

We thank Pei Zhang (Wuhan Institute of Virology) for her help with the ultracentrifugation and Electron Microscopic analysis. This work was supported by Guangdong Major Project of Basic and Applied Basic Research (2020B0301030007 to Y.S.), the Guangdong Provincial Key R&D Program (2022B1111040001 to Y.S. and W.C.), Major Project of Guangzhou National Laboratory (GZNL2023A01001 to Z.L.S), Double first-class discipline promotion project (2023B10564003 to Y.S.), and the 111 Project (D20008 to Y.S.).

## Supporting information

**Figure S1.** Necropsy of the deceased lesser panda. (A) ascites in the abdominal cavity; (B) renal necrosis; (C) hepatomegaly and steatohepatitis; (D) pulmonary hemorrhage; (E) thin and transparent intestinal walls in the small intestine; (F) mucosal injury of intestine.

**Figure S2.** Genome characteristics and sequencing depth of the LpCoV-like virus in the lesser pandas from Jiangsu (A) and Sichuan (B) provinces.

**Table S1.** Primers used for over-gap PCR.

**Table S2.** Primers used for 5’/3’RACE PCR.

**Table S3.** Universal primers used for amplify potential other segments.

**Table S4.** Detailed information of the 113 libraries.

**Table S5.** Primers used to validate subgenomic RNA.

