## Supplementary figures and images for "Identification and genomic characterization of a novel bisegmented coronavirus in the lesser panda"

### Figure S1

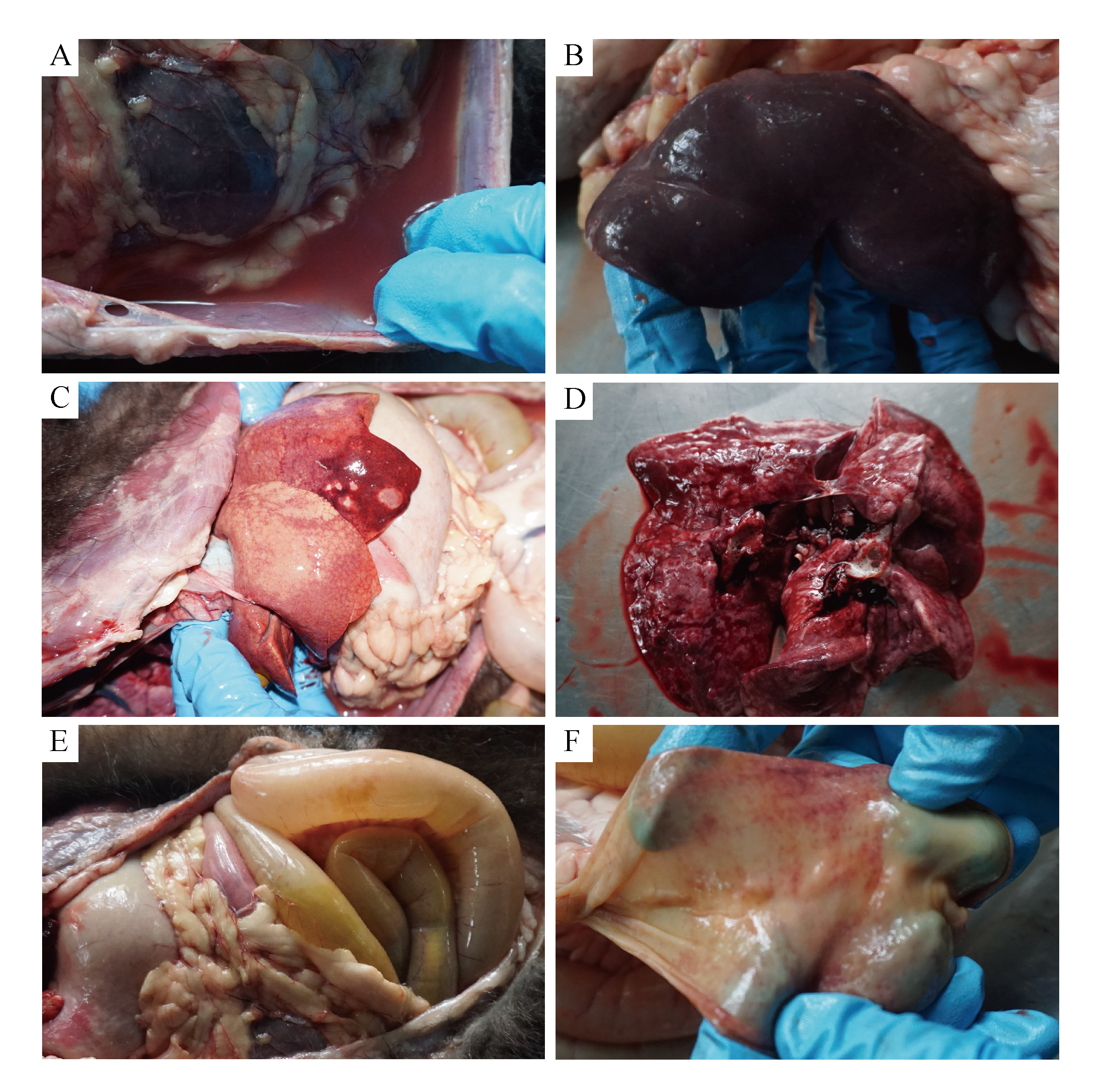

### Figure S2

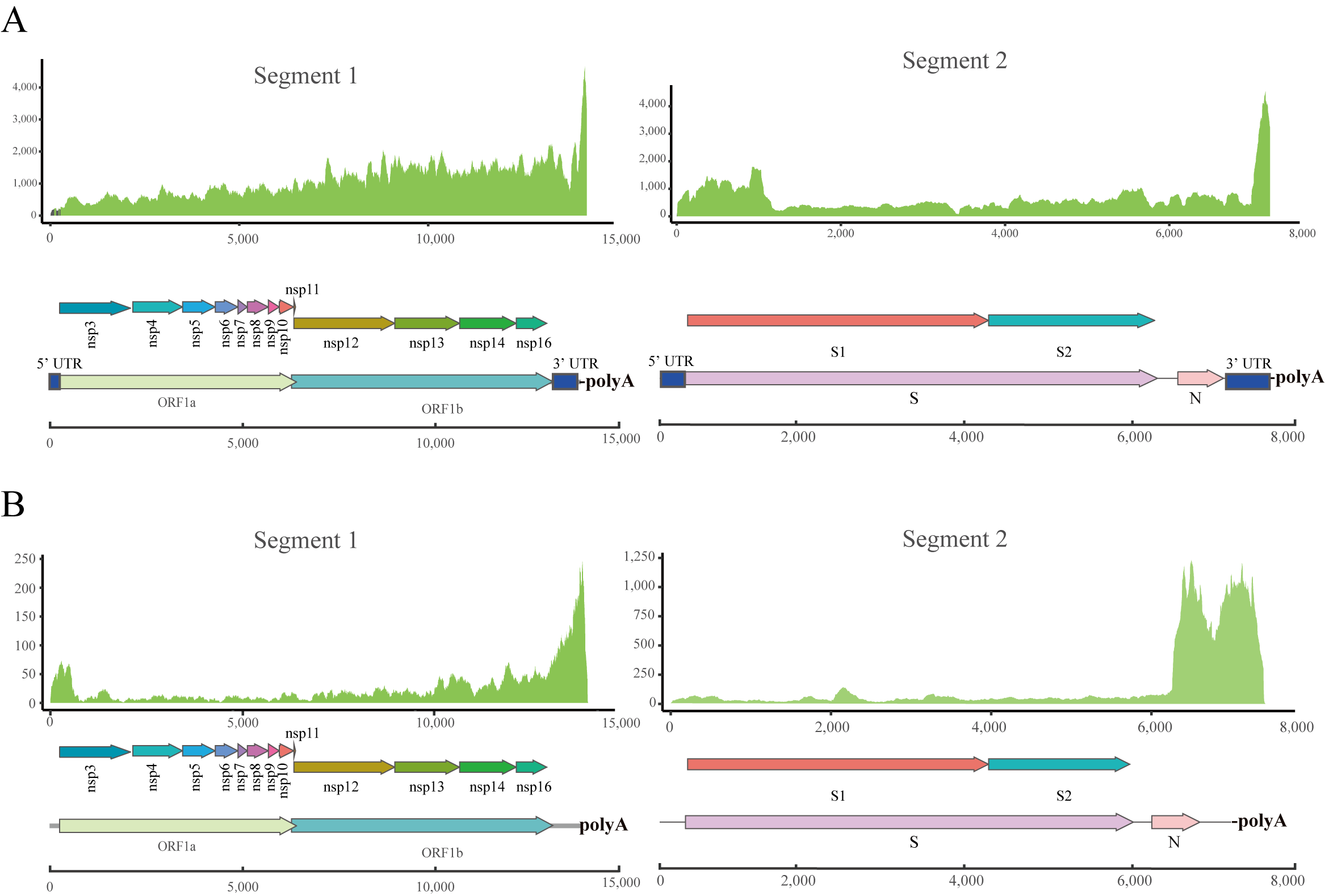
